# Identification of hub genes and pathways in nonalcoholic steatohepatitis by integrated bioinformatics analysis

**DOI:** 10.1101/2021.09.10.459743

**Authors:** Pegah Einaliyan, Ali Owfi, Mohammadamin Mahmanzar, Taha Aghajanzadeh, Morteza Hadizadeh, Ali Sharifi-Zarchi, Behzad Hatami, Hamid Asadzadeh Aghdaei, Mohammad Reza Zali, Kaveh Baghaei

**Author notes:** **Availability of data and material:** The datasets used in this paper are all available on www.ncbi.nlm.nih.gov.

## Abstract

**Background:** Currently, non-alcoholic fatty liver disease (NAFLD) is one of the most common chronic liver diseases in the world. Forecasting the short-term, up to 2025, NASH due to fibrosis is one of the leading causes of liver transplantation. Cohort studies revealed that non-alcoholic steatohepatitis (NASH) has a higher risk of fibrosis progression among NAFLD patients. Identifying differentially expressed genes helps to determine NASH pathogenic pathways, make more accurate diagnoses, and prescribe appropriate treatment.

**Methods and Results:** In this study, we found 11 NASH datasets by searching in the Gene Expression Omnibus (GEO) database. Subsequently, NASH datasets with low-quality control scores were excluded. Four datasets were analyzed with packages of R/Bioconductor. Then, all integrated genes were Imported into Cytoscape to illustrate the protein-protein interactions network. All hubs and nodes degree has been calculated to determine the hub genes with critical roles in networks.Possible correlations between expression profiles of mutual DEGs were identified employing Principal Component Analysis (PCA). Primary analyzed data were filtered based on gene expression (logFC > 1, logFC < −1) and adj-P-value (<0.05). Ultimately, among 379 DEGs, we selected the top 10 genes (MYC, JUN, EGR1, FOS, CCL2, IL1B, CXCL8, PTGS2, IL6, SERPINE1) as candidates among up and down regulated genes, and critical pathways such as IL-6, IL-17, TGF β, and TNFα were identified.

**Conclusion:** The present study suggests an important DEGs, biological processes, and critical pathways involved in the pathogenesis of NASH disease. Further investigations are needed to clarify the exact mechanisms underlying the development and progression of NASH disease.

## 1. Background

The liver is the largest internal organ involved in vital activities such as metabolizing nutrients and energy metabolism, made up of 60% hepatocyte cells and 40% non-parenchymal cells [1]. Hepatocytes deficiency in metabolite activity can accelerate non-alcoholic fatty liver disease (NAFLD). Recent studies revealed that among NAFLD patients, non□alcoholic steatohepatitis (NASH) have a higher risk of fibrosis progression [2, 3]. NASH is a significant risk factor for end-stage liver diseases such as cirrhosis and hepatocellular carcinoma (HCC). Statistical studies estimated that up to 2025, NASH-fibrosis is one of the leading causes of liver transplantation [4, 5]. The progression of NASH depends on multifactorial risk factors such as genomic variations, race, obesity, and diabetes. Moreover, many studies suggest that NASH results from several processes like abnormal lipid metabolism, oxidative stress, mitochondrial dysfunction, dysregulation of cytokines and adipokines production. However, the exact mechanism of NASH is not precisely elucidated [5–7].

Several clinical methods detect NAFLD and NASH, such as imaging modalities, blood tests, and biopsy. Also, blood tests, like serum Alanine Aminotransferase (ALT) levels, are commonly used as a surrogate marker of more severe liver status. Liver biopsy is an effective tool to distinguish between simple steatosis and nonalcoholic steatohepatitis. The hallmark of NASH pathological signs are specific cellular features such as necroinflammation, ballooned hepatocytes with or without hepatic fibrosis [2]. However, although biopsy has limitation and risk, liver biopsy remains the gold standard to validate the diagnosis [2, 7].

With the development of high-throughput techniques, which several thousand samples are tested simultaneously under given conditions, such as microarray it has become possible to analyze the samples from a proteomic, metabolic and genomic point of view to examine the pathological status of the disease much more accurately and more quickly than before. According to statistics, 20% of patients with NAFLD develop advanced NASH, and 20% of NASH patients develop cirrhosis. NASH disease and hepatic fibrosis are known as end-stage disorders, as they are often not diagnosed in early stages. Accordingly, monitoring gene expression levels by high-throughput techniques could help in research with prediction, diagnoses, prognostic and pharmacodynamics for NASH patients, or those at risk for developing NASH to cirrhosis [8, 9]. Many genomic studies in NAFLD/NASH with concern in genome expression and variations were published to determine pathogenic pathways and genomic variation profiles. Also, gathering and mining biological data at the genomic level and drawing the interaction networks like protein-protein interaction (PPI) networks have led to understanding and clarifying valuable insights on the NASH pathogenesis and identifying the role of gene expression profiles on the development of NASH. Several bioinformatics studies have demonstrated the potential hub genes, pathways, and mechanisms that play a crucial role in NAFLD progression [6, 10, 11].

In the present study, we carried out bioinformatics analyses on multiple microarray datasets, including human gene expression profiles from GEO database. Differentially expressed genes (DEGs) in NASH group were identified, and pathway enrichment analysis was performed. In order to find hub genes and GO analysis, we constructed the PPI network of DEGs. Our results suggest the vital role of JUN, IL-6, FOS, CCL2, SERPINE1, IL1B, and MYC in the pathogenesis of NASH disease.

## 2. Materials and Methods

### 2.1. Data collection

The mRNA expression datasets of NASH were searched using the keywords: ‘NASH, ‘Obesity’ and ‘Homo sapiens’[porgn: txid9606]’, and ‘Expression profiling by array’ in the Gene Expression Omnibus (GEO) database (http://www.ncbi.nlm.nih.gov/geo) [12]. After a systematic review, eleven GSE profiles were selected for primary analysis. While seven GSE profiles, including GSE83452, GSE68421, GSE24807, GSE96971, GSE49012, GSE72756 and GSE134438 were excluded from the study after quality control, four GSE profiles (GSE37031, GSE48452, GSE89632, and GSE17470) were selected for further analyses. GSE37031 was based on GPL14877 (Affymetrix Human Genome U133 Plus 2.0 Array), GSE48452 was based on GPL11532 (Affymetrix Human Gene 1.1 ST Array), GSE89632 was based on GPL14951 (Illumina HumanHT-12 WG-DASL V4.0 R2 expression bead chip), and GSE17470 was based on GPL2895 (GE Healthcare/Amersham Biosciences CodeLink Human Whole Genome Bioarray). All data were publicly available, and the present study did not involve any human or animal experimentation.

### 2.2. Identification of differentially expressed genes

To identify differentially expressed genes (DEGs), each dataset was analyzed by limma R package in Bioconductor, utilized to mine statistically significant DEGs based on the difference in their expression values between NASH and control samples [13]. Significant differential expression values were determined as a |log fold change| ≥ 1 and the adjusted P-value threshold of 0.05.

### 2.3. Integration of Microarray Data

In this study, we considered microarray datasets consist of mRNA samples for our analysis. In this regard, datasets meeting the following criteria were excluded: (1) No declaration of disease stage or shortcomings in some samples; (2) Datasets included ambiguous samples; (3) Datasets which did not include any genes with adj-P-value <0.05 and the |logFC| >1. After analyzing each dataset separately and obtaining the important genes for each dataset, we can use the robust rank aggregation (RRA) method to combine our results and have a more clean and less noisy outcome. Moreover, this method helped us gain a general idea about our result and focus better on the important genes. Therefore, we used the RRA method to aggregate up- and down-regulated gene lists after analyzing each dataset [14]. Genes with P-value <0.05 and the |logFC| >1were considered as significant.

### 2.4. Construction of Protein-Protein Interaction

For constructing a Protein-Protein Interaction (PPI) network, we inserted integrated DEGs to the STRING (http://www.string-db.org/), which is a biological database and web resource for known and predicted PPIs [15]. In order to plot the neutral interactive network, the confidence parameter was considered 0.400 (Medium Confidence).

### 2.5. Identification of Co-expression network and hub genes

To recognize the Co-expression network and hub genes, the outcome of the STRING database was reconstructed with Cytoscape software (version 3.7.0) [16]. The Molecular Complex Detection (MCODE) [17] and CytoHubba [18] plug-ins of Cytoscape were used to identify the co-expression modules and hub genes, respectively. The criteria for using MCODE were set as: Degree Cutoff = 2, Node Score Cutoff = 0.2, K-Core = 2, and Max. Depth = 100. Also, we used the Maximal Clique Centrality (MCC) method for Cytohubba.

### 2.6. Gene Set Enrichment Analysis (GSEA)

Gene Set Enrichment Analysis (GSEA) is a useful approach for interpreting gene expression data based on the functional annotation of DEGs. We can identify specific biological processes, molecular function, cellular component, and metabolic pathways by GSEA.

GSEA for DEGs was performed by Enrichr bioinformatics tool (https://maayanlab.cloud/Enrichr/) [19], to identify pathways from Kyoto Encyclopedia of Genes and Genomes (KEGG) 2019, WikiPathways 2019, and Reactome 2016, in addition to Gene Ontology (biological process (BP), molecular function (MF), and cellular component (CC)) enrichment analysis. The cut-off criterion for GSEA was adj-P-value < 0.05.

## 3. Results

### 3.1. Dataset extraction and identification of NASH-related genes

In this study, gene expression microarray data were included 101 individuals (49 healthy controls and 52 with NASH), which were obtained from 4 different datasets, summarized in Table 1. to show statistical significance (adj-P-value) versus magnitude of change (gene expression fold change) in each dataset, volcano plot was used (Fig 1). Volcano plot displaying differential expressed genes between NASH patients and healthy controls in different datasets. It plots significance versus fold-change, and by visualizing the gene expression analysis results, it helps us to better identify the genes with large fold-changes which are also statistically significant and have the potential to be biologically significant. After screening gene expression data according to the mentioned criteria (adj-P-value <0.05, |logFC| >1) from each dataset, in order to integrate collecting candidate genes from each dataset, we used RRA analysis that make a ranking list based on adj-P-value. After removing duplicate data, the obtained list contains 119 down- and 260 up-regulated genes (Table1).

**Table 1.** Characteristics of included (✓) and excluded (×) microarray datasets used for this study

| Dataset ID | Platform | Country | year | Control Samples | NASH Samples | Up-Regulated Genes Count | Down-Regulated Genes count |
| --- | --- | --- | --- | --- | --- | --- | --- |
| ✓ GSE37031 | GPL14877 | Spain | 2013 | 7 | 8 | 93 | 65 |
| ✓ GSE48452 | GPL11532 | Germany | 2016 | 14 | 18 | 32 | 24 |
| ✓ GSE89632 | GPL14951 | Canada | 2017 | 24 | 19 | 124 | 34 |
| ✓ GSE17470 | GPL2895 | USA | 2014 | 4 | 7 | 63 | 46 |
| ✗ GSE83452 | GPL16686 | France | 2019 | --- |  |  |  |
| ✗ GSE68421 | GPL16686 | Denmark | 2019 |  |  |  |  |
| ✗ GSE24807 | GPL2895 | USA | 2019 |  |  |  |  |
| ✗ GSE96971 | GPL14951 | Canada | 2019 |  |  |  |  |
| ✗ GSE49012 | GPL17470 | USA | 2013 |  |  |  |  |
| ✗ GSE72756 | GPL16956 | China | 2018 |  |  |  |  |
| ✗ GSE134438 | GPL16686 | Japan | 2019 |  |  |  |  |

**Fig 1.**
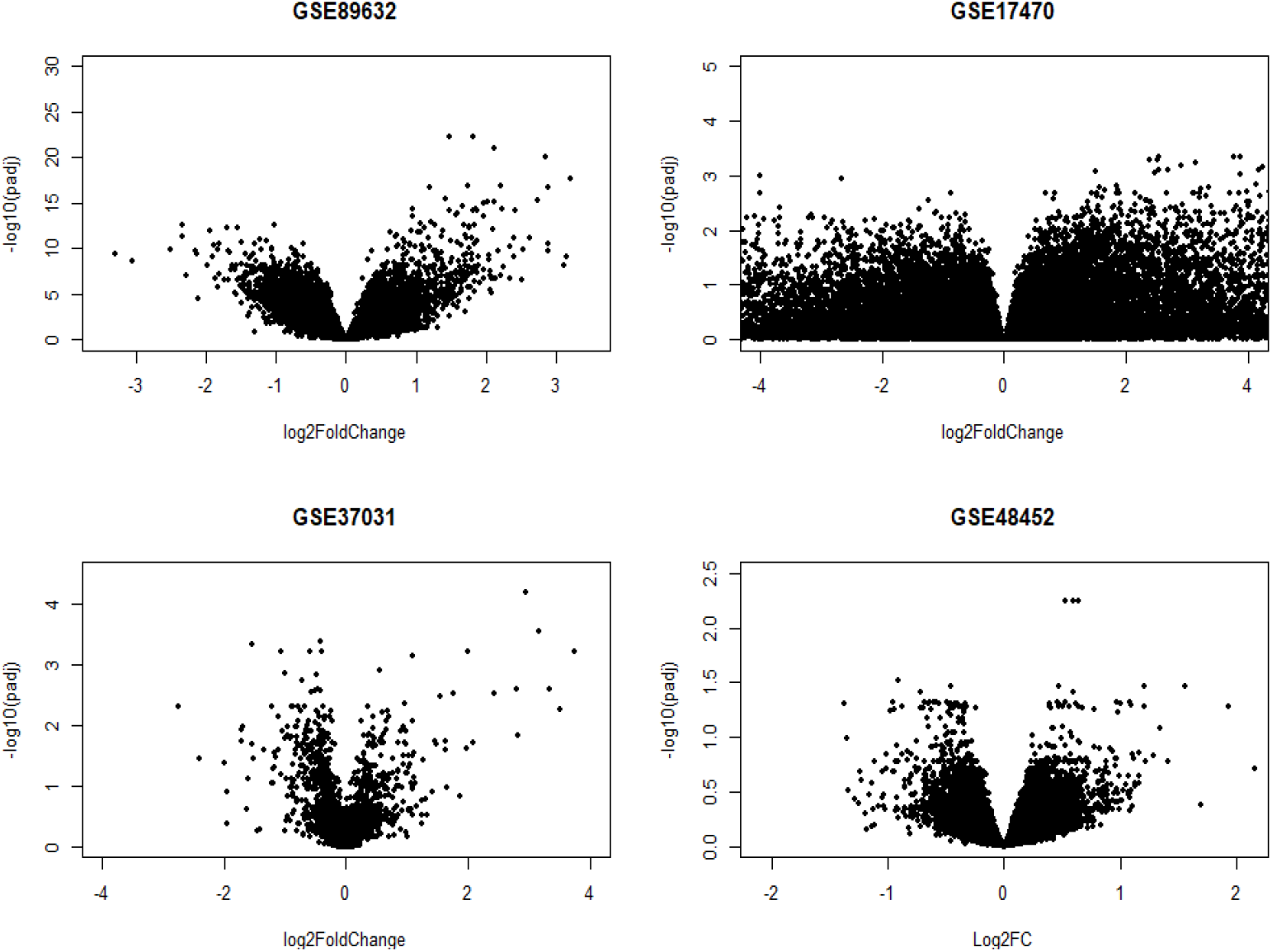
Volcano plots displaying differential expressed genes between NASH patients and healthy controls in different datasets. The vertical axis shows the expression adj-p-value of log10, and the horizontal axis indicates the log2-fold change value.

### 3.2. PPI network construction, module analysis, and selection of hub genes

To investigate the relationship between genes and their ranking in terms of the presence of this disease, a Protein-Protein Interaction (PPI) network was drawn in which each node represents a gene. Notably, genes that had disconnected from main networks were removed. Out of 379 nodes (all up- and down-regulated genes) in the network, in order to identify the hub genes, we used the Cytohubba plugin and selected the top 10 genes based on the degree and betweenness centrality measures (Fig 2). Finally, according to the highest score, we chose the top 4 modules, using MCODE (score >3). Module 1 includes 24 nodes and 110 edges with a cluster score of 9.565; Module 2, 22 nodes, 98 edges with score 9.333; Module 3, 4 nodes, 6 edges with score 4; Module 4, 16 nodes, 28 edges with score 3.733 (Fig 2).

**Fig 2.**
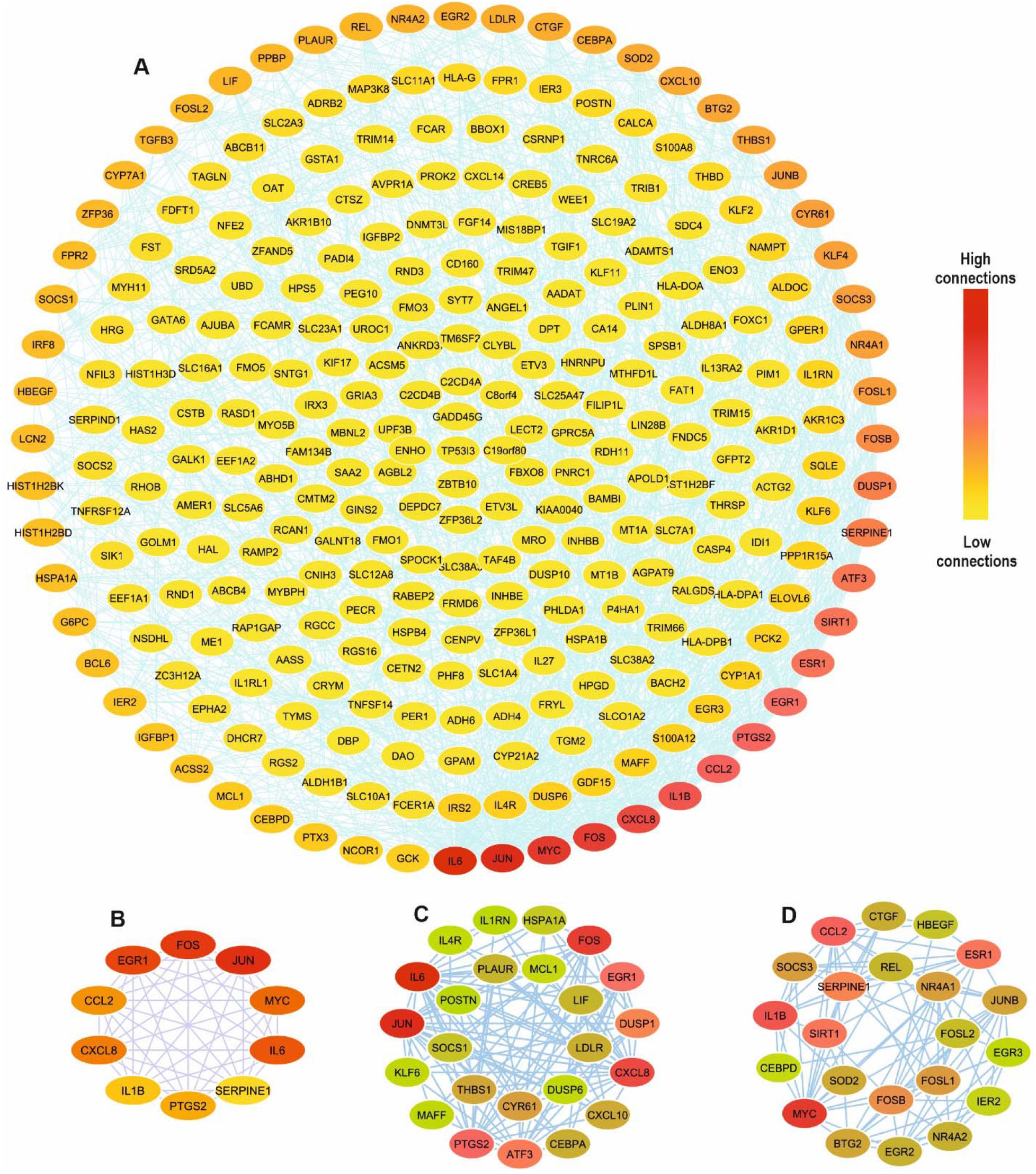
The PPI network analysis of NASH-related genes, containing GSE37031, GSE48452, GSE89632, GSE17470 datasets. (A) The overall PPI network of integrated DEGs, including 296 nodes and 1245 edges, (B) The top ten hub genes, (C) and (D) first two modules, derived from the overall PPI network. Red color indicates the high connections, and yellow color indicates the low ones.

### 3.3. Functional and pathway enrichment analyses

Due to the importance of upstream and downstream genes and possible biological pathways in the occurrence of NASH disease, pathway analysis was performed. So, three databases, such as KEGG 2019 Human, WikiPathways 2019 Human, and Reactome 2016, were used for the Pathway Analysis of Identified DEGs. The top 10 pathways related to both up- and down-regulated DEGs were summarized in Figure 3. Up-regulated genes were significantly enriched in cytokine-related pathways, including TGF-β, IL17, TNF, and oncostatin M, as well as Nuclear receptors, vitamin D and vitamin 12 pathways (Fig 3A). Down-regulated genes were enriched mostly in Gluconeogenesis, cholesterol biosynthesis, and gene regulation by SREBF (Fig 3B).

**Fig 3.**
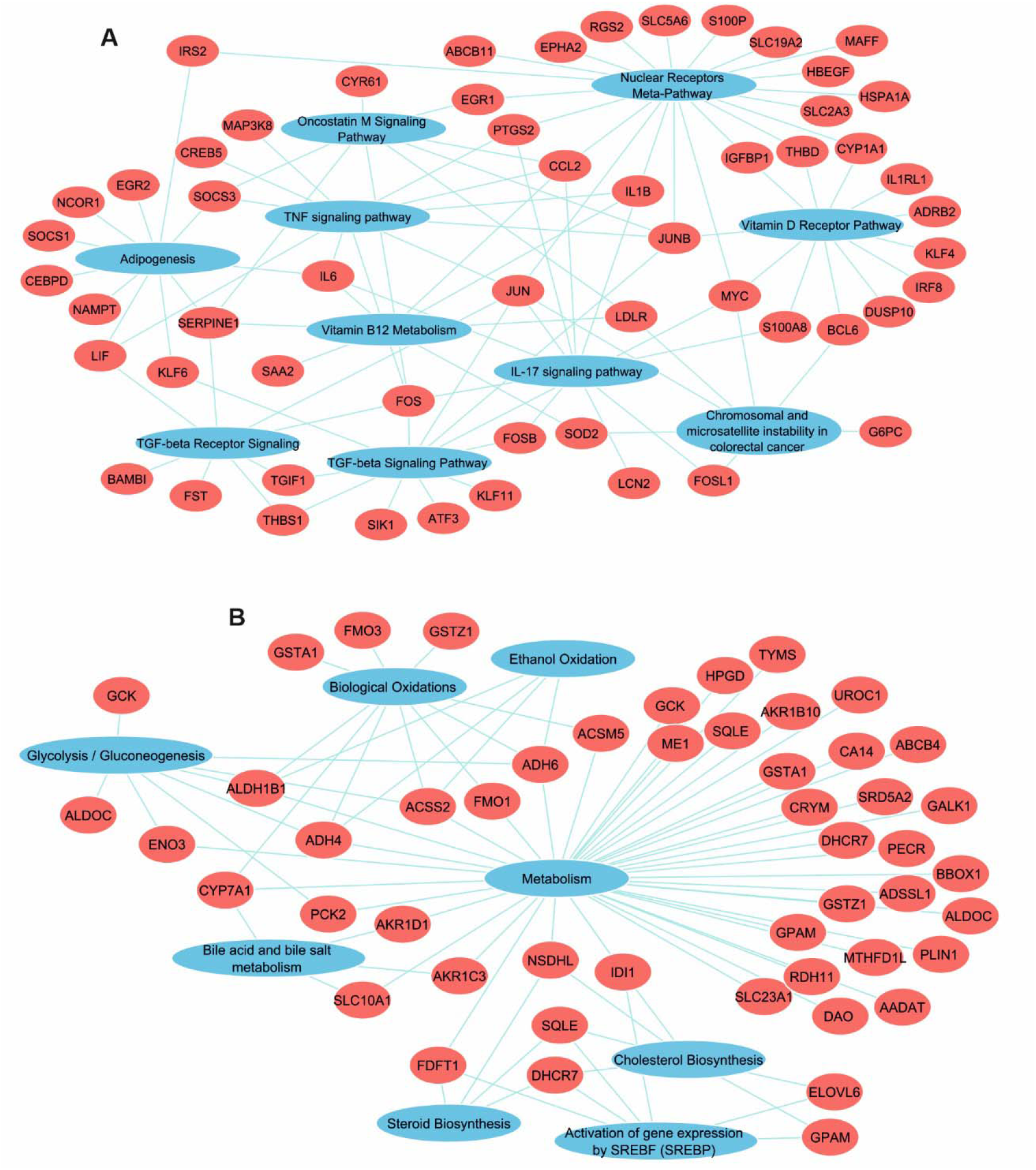
Pathway enrichment analysis for up-(A) and down-regulated (B) genes, using KEGG, Reactome, and Wikipathways databases

According to the datasets analyzed and the hub genes extracted, by considering the adj-P-value < 0.05, we obtained the top 10 Biological Process and Molecular Functions which were summarized in Table 2. No Cellular component term was significant in our GO analysis for hub genes. For BP, hub genes were mostly enriched in cytokine-mediated terms, inflammatory processes, response to lipopolysaccharide, and regulating DNA transcription. In MF analysis, hub genes were mainly associated with cytokine and chemokine activity, transcription factor and DNA binding, RNA polymerase II activity, and R-SMAD binding (Table 2).

**Table 2.**
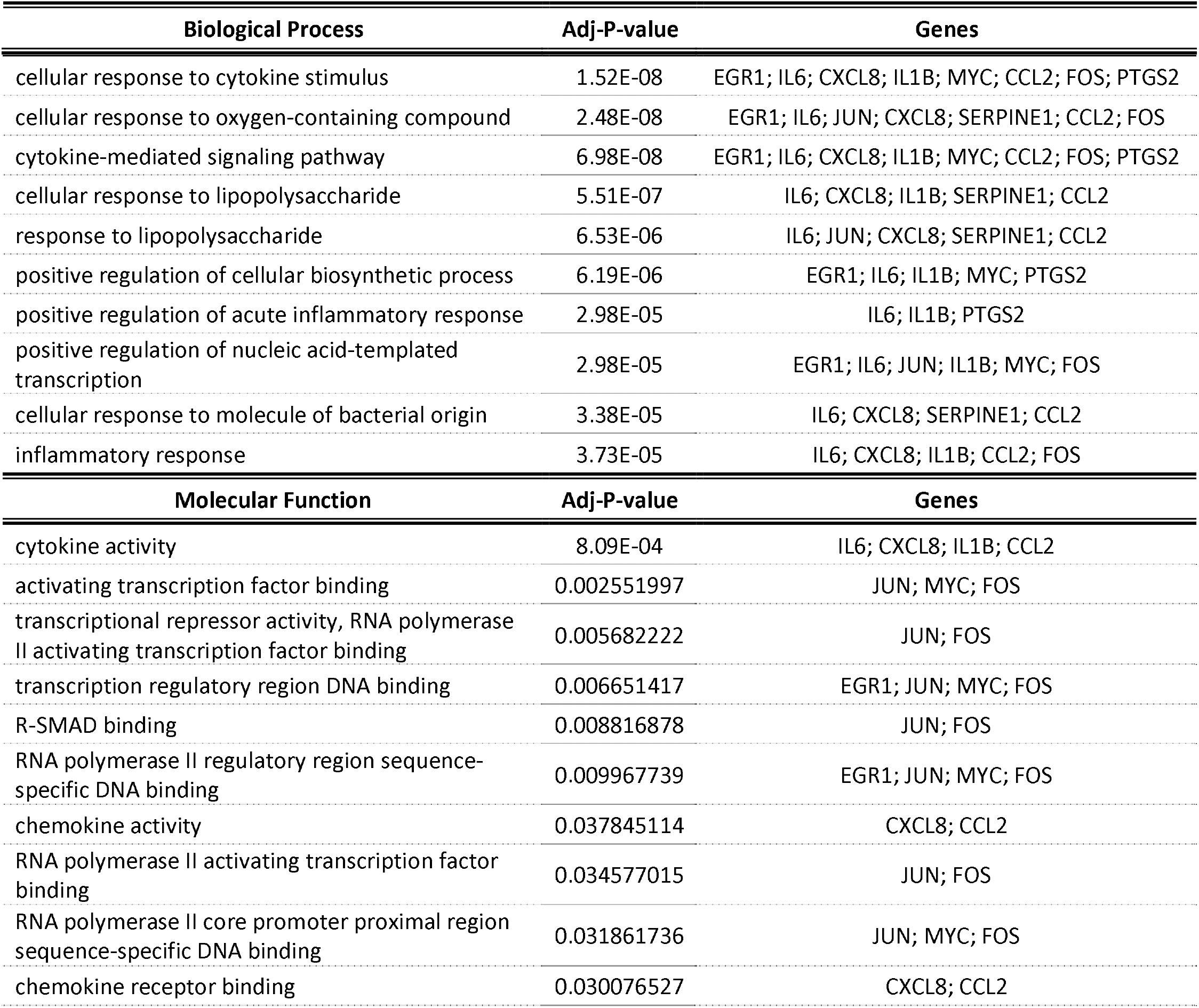
Top 10 GO enrichment analysis for hub genes. No terms were significantly enriched in the ‘cellular component’.

## 4. Discussion

Non-alcoholic fatty liver disease (NAFLD) is the most common chronic liver disease, ranging from simple steatosis (fatty liver) to non-alcoholic steatohepatitis (NASH), hepatic fibrosis, cirrhosis, and liver cancer (HCC) [20]. NASH is the aggressive form of NAFLD, characterized by steatosis, cell injury, inflammation, and hepatocyte ballooning [21]. Although liver biopsy is the gold standard to diagnose NASH and fibrosis, it has some limitations and risks which prevent it from being used in the early stages of the disease. Therefore, exploring the molecular mechanisms of NASH is pivotal to improve early diagnosis.

In the present study, we used integrated bioinformatic methods for transcriptomic analysis. So, four microarray datasets (GSE89632, GSE37031, GSE17470, GSE48452) were analyzed, and a total of 379 DEGs, including 260 up- and 119 down-regulated genes, were identified and integrated by the RRA algorithm. Using STRING and Cytoscape, we constructed a PPI network for DEGs, and the MCODE plugin performed cluster analysis. Finally, 10 hub genes associated with NASH were obtained via the Cytohubba plugin, including JUN, IL-6, FOS, EGR1, CCL2, SERPINE1, CXCL8, PTGS2, IL1B, and MYC. All hub genes were among up-regulated DEGs and demonstrated the importance of these genes.

Feng et al. proved the significance of FOS as a hub gene in NAFLD/NASH diseases. It is demonstrated that CAT, ALDH2, and ALDH8A1 are associated with NASH progression to HCC [6]. A recent bioinformatics study has shown that IL-6, MYC, FOS, FOSB, and JUNB are among NASH hub genes, as well as IL-17 and TNF-α signaling pathways were enriched in KEGG analysis [11].

Our pathway analysis for up-regulated DEGs revealed that they are enriched in immune system pathways such as TGF-β, TNF-α, IL-17, and oncostatin M signaling pathways. TGF-β signaling regulates a wide range of cellular processes, such as proliferation, differentiation, migration, apoptosis, and specification, by binding to the serine/threonine kinases receptors on the cell surface [22]. It has been shown that increased TGF□β signaling in hepatocytes contributes to lipid accumulation by regulation of lipogenesis and β□oxidation□related gene expression. Also, TGF□β Induces cell death in steatotic hepatocytes through Smad and Rho/ROCK Signaling pathway, resulting in inflammation and fibrosis [23].

Our analysis demonstrated that FOS (c-Fos) is a crucial gene in the NASH and is involved in the signaling pathways, including TGF□β, IL-17, TNF-α, and oncostatin M. It has been proved that TGF-β stimulates c-Fos, and c-Fos expression in mice liver increases immune cell infiltration and inflammation [24, 25]. Also, TNF-α-which is an essential mediator of inflammation- production will be elevated in LPS injected Fos−/− mice. So, FOS might act as an inflammatory response suppressor [26]. TNF-α is overexpressed in the liver and adipose tissues of obese patients, and the expression of TNF-α receptor-type1 mRNA is higher in NASH patients’ liver tissue [27]. Manco et al. researched on the serum levels of TNF-α in children with NAFLD and showed that TNF-α levels were significantly associated with a NAS score of 5 and more [28]. IL-17 and TNF-α pathways were also in PPI clusters 1 and 2, where IL-6 was enriched in both of them. Many studies have been reported the elevated levels of IL-6 and TNF-α expression in blood and serum samples of NAFLD patients, correlated with steatosis and NASH progression [29–31]. IL-6-deficient displayed less lobular inflammation and serum ALT than wild-type in MCD diet-induced NASH mice [32]. Conversely, some studies reported that the blocked IL-6 signaling or IL-6-deficient in MCD or HFD diet-fed mice results in steatosis and inflammation, demonstrating the protective role of IL-6 against the NASH development [33, 34]. Association of IL-6 with TGF□β promotes Th17 expression, which produces IL-17, a pro-inflammatory cytokine, inducing the production of other cytokines such as TGF□β and TNF-α. Therefore, it has been implicated in liver inflammation and fibrosis progression by activation of Kupffer cells. It has been reported that IL□17 mRNA expression was increased in NASH patients compared with healthy controls and plays a pivotal role in hepatocyte steatosis and hepatic inflammation associated with NASH [35]. Besides, it has a fibrogenic effect by stimulating Kupffer cells to express cytokines, like IL-6, IL-1, TNF- α, and TGF□β, and stimulating hepatic stellate cells to express collagen-1 and transformation to myofibroblasts by Stat3 [36]. There are some reports about the relation between IL-17 and adipogenesis. It suppresses adipogenesis by inhibiting expression of several pro-adipogenic transcription factors, including PPAR-γ, C/EBPα, and KLF15, and enhances the expression of anti-adipogenic TFs such as KLF2 and KLF3 [37]. PPAR-γ and C/EBPα are the central regulators of lipogenesis, which induces adipocyte-specific genes, including fatty acid-binding protein 4 (FABP4), Glut4, and phosphoenolpyruvate carboxykinase (PEPCK) [38]. Also, PPAR-γ, which is elevated in the fatty liver, inhibits HSCs activation and fibrogenesis, and its deficiency protects against steatosis [39]. Vitamin D regulates the metabolism of free fatty acids by its effect on PPAR-γ and relieving FFA-induced insulin resistance in vitro. Many studies have been found that 25(OH)D has lower levels in NAFLD/NASH patients than healthy controls [40]. Ding et al. reported that treatment of CCL4-induced mice with calcipotriol, a vitamin D receptor agonist, decreased fibrotic marker expressions such as COL1A1, TGF-β1, and TIMP-1 [41]. Additionally, adipose tissue can directly affect hepatic cells. Induction of visceral adipose tissue-derived exosomes into HepG2 and HHSteC cells increased TIMP-1 and integrin αvβ5 and decreased MMP-7 and PAI-1 expression in HepG2 cells, and increased TIMP-1, TIMP-4, Smad-3, integrins αvβ5 and αvβ8, and MMP-9 expressions in HSCs. The effect of integrin αvβ5 on fibrosis induction in other organs through the TGF□β pathway suggests the same function in the pathogenesis of NAFLD [42]. In addition to the critical role of IL-6 by itself, Oncostatin M (OSM), a member of the IL-6 cytokine family, synthesized by activated monocytes and macrophages, activated T cells, dendritic cells, neutrophils, and eosinophils. Some investigations show that OSM receptor β increases in the hepatocytes of the mouse model of hepatic steatosis. OSMRβ-deficient mice exhibit delayed hepatocyte proliferation and persistent liver necrosis after CCl4 exposure, resulting in elevation of MMP-9, suppression of TIMP□1 and TIMP□2, and matrix degradation. So, CCl4-induced liver damage worsens in OSMRβ−/− mice. Additionally, OSM is expressed much more during NASH than that during steatosis in mice [43, 44]. A recent study revealed that OSM expression has increased in human liver specimens from NAFLD patients, especially in the NASH stage, and there was no up-regulation in simple steatosis [45]. Up-regulated genes were also enriched in vitamin B12 metabolism. A recent study showed that serum levels of B12 have a negative correlation with NASH severity, whereas another research reported no difference in serum vitamin B12 levels between NAFLD/NASH patients and healthy controls [46–48]. As a result, more studies are needed to investigate this issue.

In the case of down-regulated genes, the pathway analysis was mainly enriched in gluconeogenesis and steroids biosynthesis, such as cholesterol and gene expression by SREBF. Liver steatosis induction by HFD diet in mice resulted in a significant increase of liver and plasma free fatty acid and triglyceride levels and plasma ALT activities. Moreover, gene expression analysis reveals that genes encoding cholesterol biosynthesis enzymes, such as squalene epoxidase (SQLE) and NAD(P)-dependent steroid dehydrogenase-like (NSDHL) were down-regulated [49]. These results confirm our analysis, in which SQLE and NSDHL were enriched in the cholesterol, sterol, and SREBF pathways. Liu et al. performed RNA-seq analysis of NAFLD-HCC tumor tissues and showed that SQLE is overexpressed in NAFLD–induced HCC patients. Further investigations on *Sqle* transgenic mice demonstrated higher hepatocyte ballooning, inflammatory cell infiltration, and serum ALT and AST levels than wild type. Also, proinflammatory mediators mRNAs, including CCL-12, CCL-20, CCR-5, CXCL9, CXCR4, IL-2, IL-4, IL-18r1, OSM, and SPP1 were increased [50]. So, more studies are needed to find out the exact mechanism of SQLE in NASH. A recent study on the Hela cell line showed that SQLE up-regulation is regulated by SP1 and SREBF2 transcription factors, increasing cholesterol and cholesteryl ester accumulation [51].

We performed GO analysis for hub genes, and the results indicated that most of them are involved in biological processes such as ‘cellular response to cytokine stimulus’, ‘cytokine-mediated signaling pathway’, and ‘inflammatory response via cytokine and chemokine activity’ molecular functions. A new meta-analysis study revealed that the concentration of CCL2 and CXCL8 in the NAFL and CCL3, CCL4, CCL20, CXCL8, and CXCL10 in the NASH patients was higher than that in the control group, respectively. It has been found that CCl2 is secreted by activated HSCs and Kupffer cells, which contributed to the lipid accumulation in hepatocytes and increased macrophage accumulation in the liver and adipose tissue, leading to fatty liver. Also, activated HSCs and Kupffer cells secrete CXCL8, which has an important role in NASH by recruitment of neutrophils via activation of the AKT/mTOR/STAT3 pathway [52]. An in vitro study showed that TGF-β1 significantly up regulated CCL2 and CXCL8 concentrations in HSCs cell line [53]. Gore et al. studied on the inflammation in an ex vivo NASH murine model and reported that the gene expression of IL-6, IL1B, and TNF-α was increased [54]. Interestingly, IL1B in association with IL-23 induced IL-17, IL-21, and IL-22 expression by gamma delta (γδ) T cells in the absence of additional signals [55]. JUN and FOS are AP-1 transcription factor subunits, enriched in activating transcription factor binding molecular function. Some studies observed an increased level of c-Jun-the best-characterized AP1 component- in mouse models and human NAFLD patients, showing a correlation between the inflammatory level and hepatocytic c-Jun expression [56, 57]. Expression of c-Fos in hepatocytes leads to liver inflammation so that granulocytes forming the bulk of recruited immune cells. Also, c-Fos influences the LXR/RXR pathway and controlling hepatic cholesterol, which its high expression leads to reduced pathway activity and accumulation of total and free cholesterols. It has been reported that PPAR-γ is responsible for decrease LXR/RXR pathway activity [25]. Our GO analysis revealed that MYC (c-Myc) performs many biological processes, especially in ‘inflammatory pathways’, and interacts with many other inflammation mediators such as IL-1, 2, 4, 8, 10, TNF-α, NF-κB, PPAR-Y, and p53. IL-6-c-Myc and IL-6-AP-1-c-Myc pathways increase c-Myc expression, whereas IL-6-NF-κB-c-Myc pathway inhibits that. TGF-β increases c-Myc expression via stimulating Smad3 and downregulates it by the Smad2 [58]. SERPINE1, also known as PAI-1, was enriched in ‘response to lipopolysaccharide’ BP. PAI-1 is a downstream target of TGF-β, assessed as an antithrombotic agent, and its plasma concentration is correlated with the increasing NASH/fibrosis stage [59, 60]. Augmented levels of PAI-1 have been considered as a risk factor for thrombosis and cardiovascular events by inhibiting tissue□type plasminogen activator (tPA) and urokinase plasminogen activator (uPA), the main enzymes for inducing fibrinolysis, results in fibrin accumulation [61]. Furthermore, this process occurs in alcohol□induced liver injury, in which ethanol causes PAI-1 activation, leading to fibrin accumulation in the liver by inhibiting its degradation. Fibrin deposition sensitizes the liver to LPS□induced necrosis and inflammation. [61, 62]. These findings link NAFLD/NASH to cardiovascular disease (CVD) and might elucidate the increased CVD risk associated with NAFLD.

## 5. Conclusion

In conclusion, by performing transcriptomic analyses on NASH datasets, our study illustrated the potential signaling pathways and processes that may play a critical role in the molecular pathogenesis of the disease. Our results suggest the crucial functions of inflammatory responses and adipogenesis through the significant role of TGF-β, TNF-α, and IL-17, as well as the association of hub genes such as JUN, IL-6, FOS, CCL2, SERPINE1, IL1B, and MYC. These findings may serve as useful markers for future studies on NASH diagnostic and therapeutic.

## Acknowledgments

The authors would like to thank the Research Institute of Gastroenterology and Liver Diseases of the Shahid Beheshti Medical University for its financial support of this study

## Abbreviation

NAFLD: Non-alcoholic fatty liver disease
NASH: non-alcoholic steatohepatitis
GEO: Gene Expression Omnibus
DEGs: Differentially Expressed Genes
RRA: Robust Rank Aggregation
PPI: Protein-Protein Interaction
GSEA: Gene Set Enrichment Analysis
KEGG: Kyoto Encyclopedia of Genes and Genomes
BP: Biological Process
MF: Molecular Function
CC: Cellular Component

---

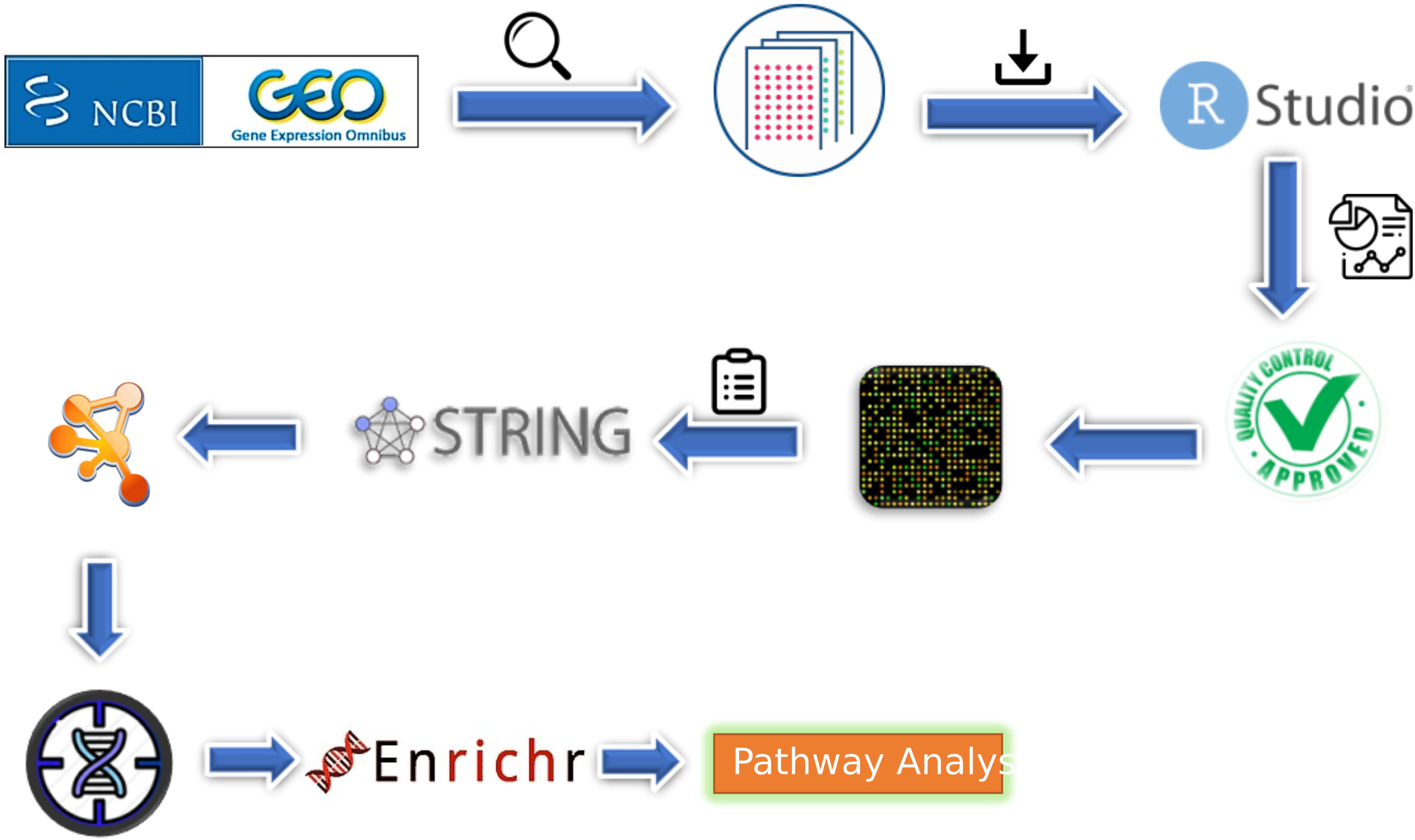

